# Fate Bias and Transcriptional Memory of human B cells

**DOI:** 10.1101/2022.07.14.499766

**Authors:** Michael Swift, Felix Horns, Stephen R. Quake

## Abstract

Lineage tracking offers a direct approach to study cell fate determination. In this work we combined single cell transcriptomics and lineage tracing to better understand fate-choice in human B cells. Using the antibody sequence to trace cell lineage during *in vitro* differentiation, we identified intrinsic proliferative and cell fate biases of B cell subtypes. Clonal analysis revealed that IgM memory B cells were more proliferative than any other B cell subtype, and that cells from the same clone had highly concordant fates. We found that transcriptional memory within clones varies across genes, with strongest persistence in genes related to cell fate determination. Similar persistent transcriptional programs were observed in human plasma cells from bone marrow, suggesting that these programs maintain long-term cell fate in vivo. These results show that cell-intrinsic fate bias influences human B cell differentiation and reveal molecular programs underpinning cell fate determination in B cells.

## Introduction

A key focus of developmental biology is the relationship between the molecular milieu of a progenitor cell and its differentiation outcomes. These outcomes are variously referred to as cell fate, cell identity, or cell state. Lineage tracing offers a powerful way to map which progenitor cells adopt which cell fates. Even rudimentary cell labeling techniques show clonally related offspring are biased towards similar cell fates (Whitman, 1878), and recent technological advances confirm the same with greater throughput and resolution. However, the contribution of cell-extrinsic versus cell-intrinsic molecular factors as determinants of cell fates remains largely unknown.

In order to better understand cell fate determination, multiple groups have used high-throughput sequencing to measure endogenous or transgenic DNA barcodes as labels of cellular lineage (Lu et al. 2011; Horns et al. 2016; Ludwig et al. 2019). Recently, it is possible to use high-throughput sequencing to perform both lineage tracing and transcriptomics in single cells. This combination directly measures the molecular relationships between proginitors and their offspring, allowing stronger inference of molecular determinants of cell fates (Biddy et al. 2018; Weinreb et al. 2020). For example, by analyzing the transcriptomes of lineages biased towards efficient reprogramming outcomes, Biddy *et al*. were able to identify a previously uncharacterized gene which increased stem-cell reprogramming efficiency by three-fold.

In the human immune system, clonal lineages of leukocytes rapidly proliferate whilst adopting diverse cell fates. This dynamic occurs *in vivo* as a response to complex pathogenic challenges such as viruses, bacteria or cancer. Spatially organized cellular structures, called germinal centers, orchestrate this process *in vivo*. However, *in vitro* differentiation protocols using only T or B cells can recapitulate important features of the germinal center, and provide valuable insight into the process (Tangye et al. 2003; Hasbold et al. 1998). *In vitro*, a researcher can control most extrinsic factors, such as cell density, cytokine cocktails, and media compositions, allowing them to study cell-intrinsic differentiation programs. In the case of B and T cells, (Cheon et al. 2021) well-controlled extrinsic conditions still reliably generate a large diversity of cell fates, indicating a strong contribution of intrinsic cell diversity to population-level diversity seen *in vivo*. The question of how gene expression responds to extracellular stimuli, while a cell maintains its erstwhile identity is generally poorly understood. And the transcriptional programs underlying cell-intrinsic clonal fate bias remain unknown. Furthermore, how extrinsic signals, intrinsic state, and clonal population structure interact to determine the dynamics of the B cell immune response remains poorly understood.

Here, we gain insight into some of these gaps in knowledge by obtaining lineage and single-cell RNA sequencing measurements of differentiating human B cells. We used the B cell receptor (BCR) gene as a genetic lineage marker and we paired this information with a transcriptomic readout of cellular identity during an *in vitro* differentiation of human B cells. These data allow us to quantitatively infer the intrinsic biases B cells have towards specific cell fates, analyze the clonal dynamics of *in vitro* B cell activation, and identify transcriptional memory states both *in vitro* and *in vivo*.

## Results

### *In vitro* B cells recapitulate major aspects of *in vivo* B cell development

To gain insight into differentiation and fate choice of human B cells, we performed an *in vitro* differentiation protocol using primary human B cells from healthy donors. We stimulated the B cells with cytokines to induce proliferation, class-switch recombination (CSR), and reprogramming into terminally differentiated Plasma B cells. We performed single-cell RNA-sequencing on time-course samples from this protocol (Fig 1A), which furnished us with genetic and transcriptional information about B cell differentiation. Additionally, we contextualized our *in vitro* differentiation by (1) integrating our single cell RNA sequencing data with publicly available data from 10X Genomics and (2) measuring CD138+ bone-marrow plasma cells from a separate donor (i.e. terminally differentiated B cells *in vivo)*. After low quality and non-B cells (Fig S1A & B), we obtained the mRNA transcriptome and VDJ sequences for 29,703 B cells (Fig 1C).

**Figure 1.**
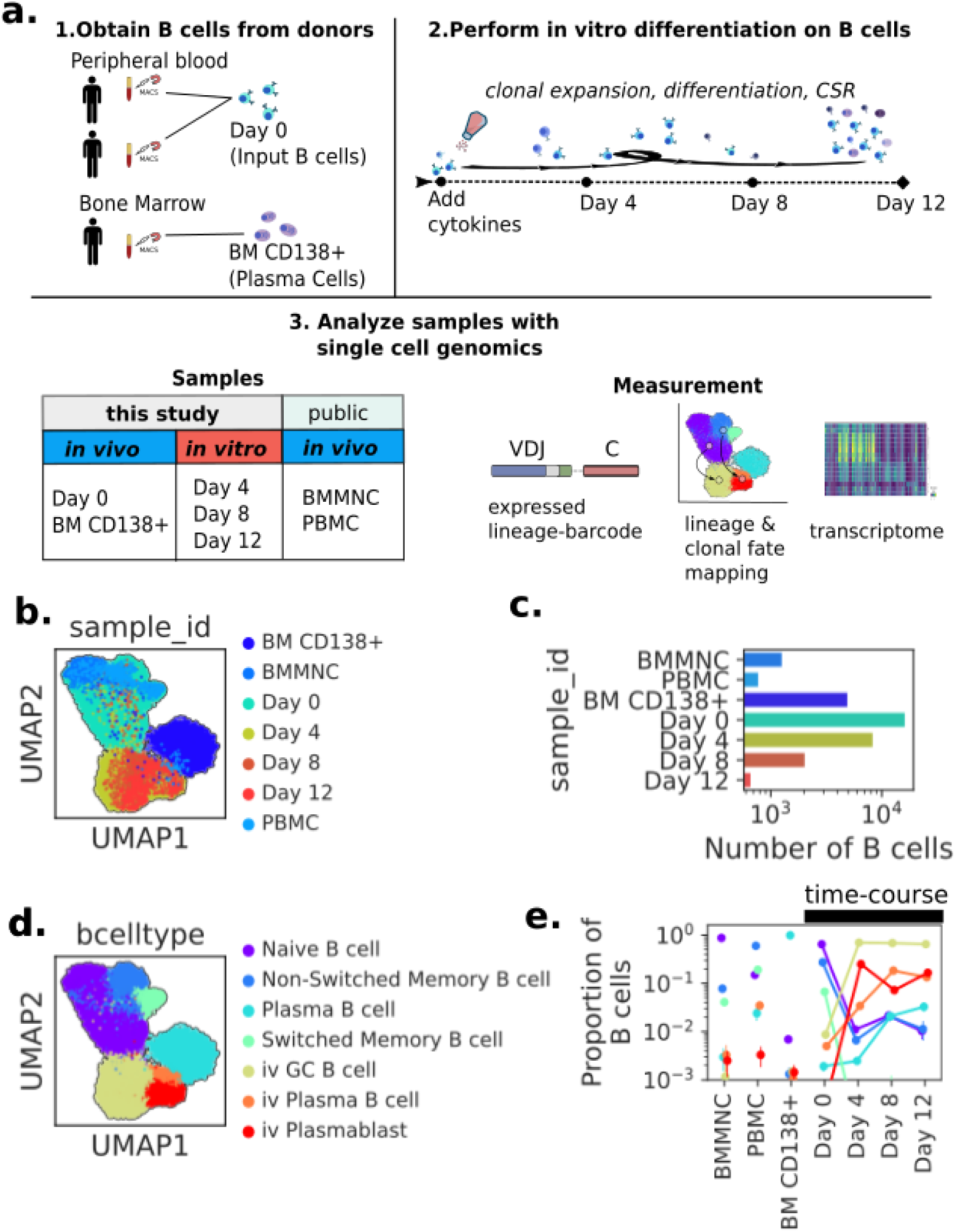
Experimental Overview for studying in vitro B cell Dynamics using integrated single cell genomics and lineage tracing. (A) Experimental Overview. (1) B cells or Plasma cells were purified from blood or bone marrow respectively using MACS. (2) B cells from the same purification were stimulated with the StemCell B cell expansion kit. Samples of the *in vitro* differentiation were collected on Day 4, 8, and 12. (3) Schematic of the single cell genomic data collected and analyzed. (B) Principal-component analysis and UMAP embedding separates cells into distinct clusters. Each dot is a cell, colored by sample of origin and the countplot indicates the number of cells passing QC from each sample. (C) Countplot of the numbers of B cells passing QC for each sample. (D) UMAP embedding with cell type annotations; see S1F for celltype defining genes. (E) Proportion of b cell types in each sample. Samples from the time course are connected. All error bars are 95% confidence intervals calculated by resampling.

**Figure 2.**
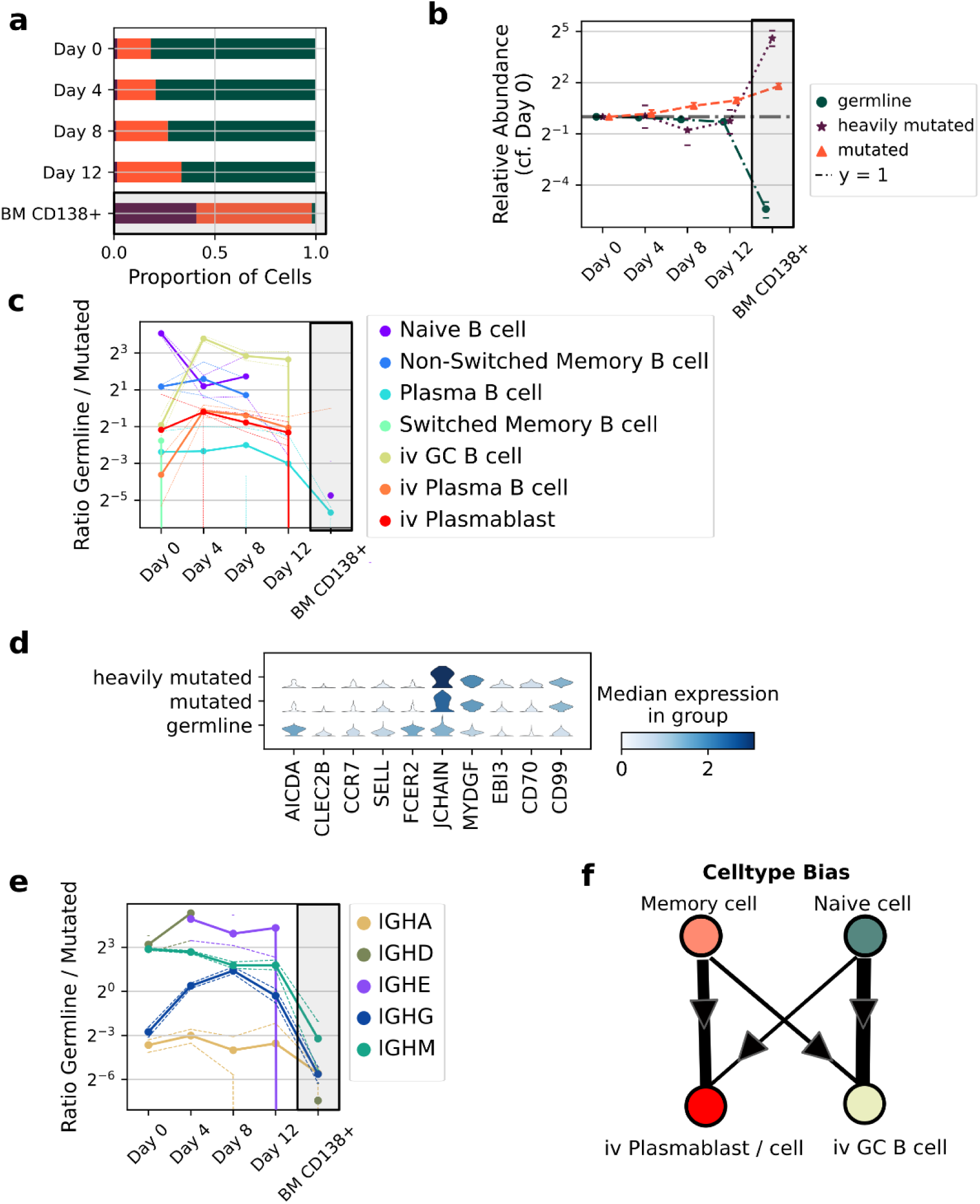
Characterization of cell intrinsic phenotypes using VDJ mutation status. (A) Proportion and (B) relative abundances (right) of germline, mutated, and heavily mutated B cells in the samples. Error bars in all figures are 95% confidence intervals calculated by resampling; gray boxes throughout the figure indicate BM CD138+ is not part of the time-course but serves as a comparison sample (C) Ratio of germline to mutated cells in each cell state over the time-course. (D) Differential gene expression in Day 4-12 B cells between the mutation levels. Gene expression is log base 2 umis per 10,000. (E) Ratio of germline to mutated cells for each isotype group. (F) Illustration of the inferred celltype biases.

**Figure 3.**
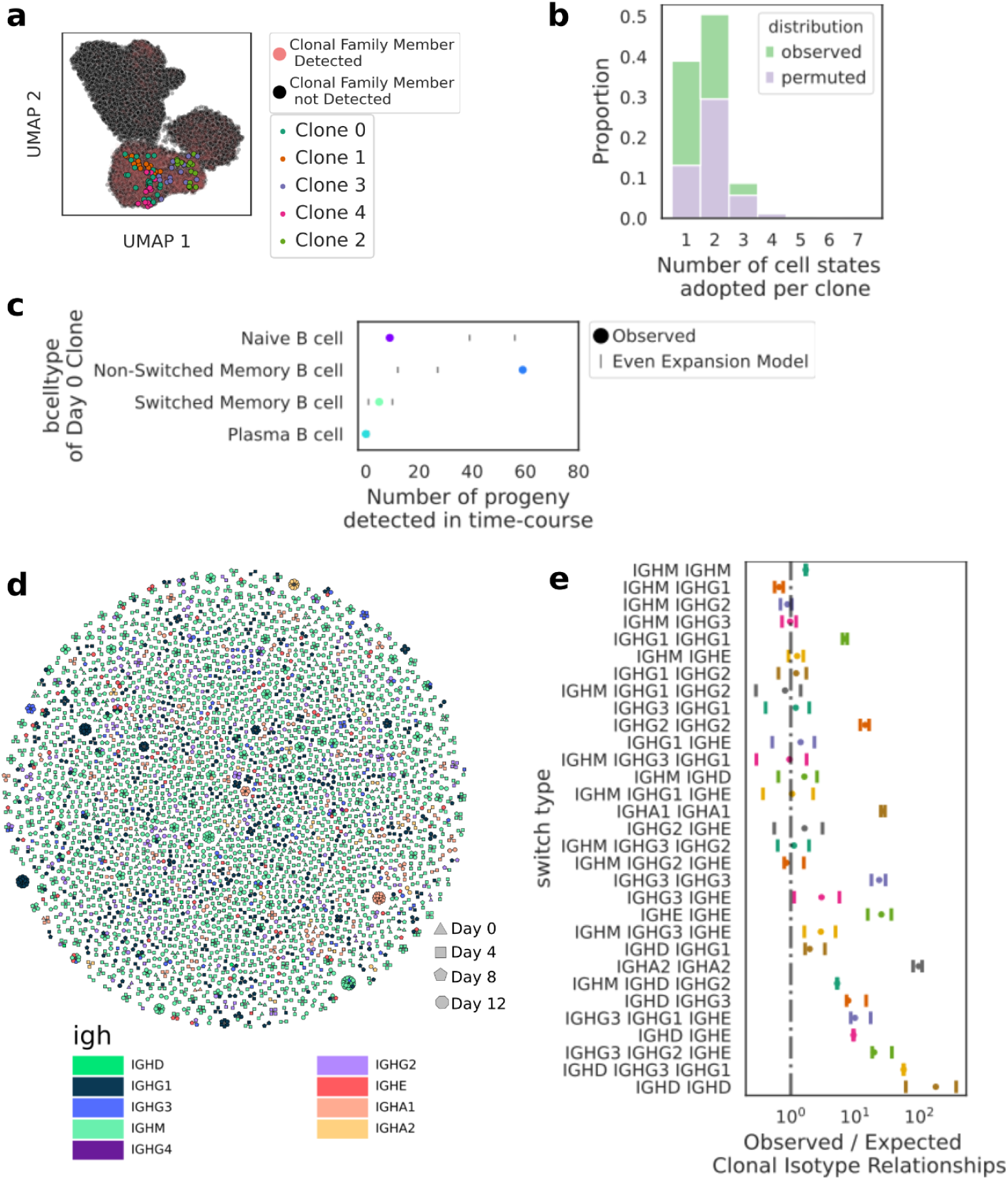
Clonal families allow inference of intrinsic proliferative ability, limited clonal fate outcomes, map of class-switching *in vitro*. Error bars in all figures are 95% confidence intervals calculated by resampling (A) UMAP embedding showing the largest clonal families detected (B) Stacked histogram showing the number of cell fate outcomes available within clones compared to permuted clonal labels (C) Progeny of Non-switched Memory B cells are overrepresented in the differentiated population. Observed data are compared to a model where the observed Day 0 clonal structure is expanded evenly for 8 divisions and then randomly resampled (D) A graph-based representation of clonal families. For each cell (node), the isotype (color), time-point (shape). (E) Ratio of observed to expected clonal isotype relationships for clonal isotype relationships that were detected.

**Figure 4.**
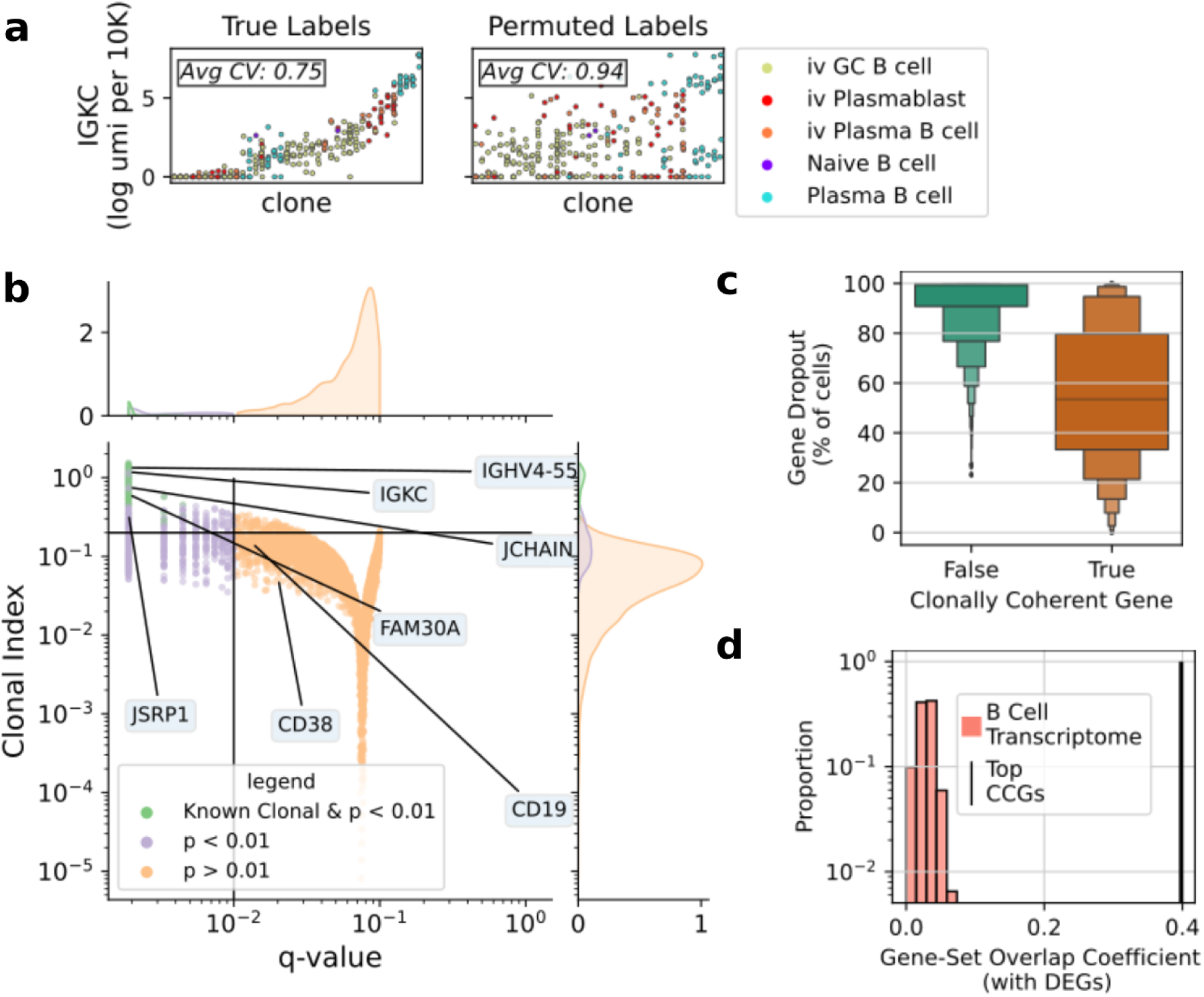
Clonal transcriptional programs are strongly enriched for fate determining genes. (A) Cascade plots where each column is a clonal family of size >=5. Families are rank-ordered by the mean gene expression of the family. Each dot is a cell, colored by the cell state label. The true clonal families are plotted on the left and a permutation of the clonal labels is plotted on the right. (B) A volcano plot showing results of the transcriptome-wide permutation test. Q-values are the Benjamini-Hochberg corrected p-values and the clonal index is a normalized metric of expression variance described in the methods. Genes of interest are labeled and groups of interest are colored. (C) Clonal genes were likely to be detected in many cells. Boxen plot showing the distribution of percent dropout for genes tested (D) The set of top CCGs strongly overlap with the set of top cell-state defining genes compared to sets of randomly selected genes (p < 0.001). The null expectation is the B cell Transcriptome: randomly sampled sets of genes expressed in at least 20% of B cells.

Dimensionality reduction by principal component analysis (PCA) and UMAP (McInnes et al. 2018) of the single cell transcriptomes revealed several distinct cell states, which we annotated based on known markers (1D, S1E). We analyzed the relative abundances of these cell states over time, and found that they changed dramatically (1E). First, we noted that non-B cells present in the input rapidly became undetectable by day 4, which shows the specificity of the cytokines for B cell expansion. Other notable dynamics included a three-fold decrease in the relative abundance of plasma cells from Day 0 to Day 4 in favor of an increase in cycling, germinal center-like B cells. However, in Days 8 and 12, we observed an increased abundance of plasma cells indicating the differentiation of resting Naive and Memory B cells types into Plasma cells. Finally, we note the majority of cells began to die in culture by Day 10, suggesting cell death was a major contribution to the population dynamics by Day 12.

### Lineage tracking of VDJ mutation status allows inference of cell-fate biases

Upon experiencing antigenic stimulation, Naive B cells genetically diversify their BCR by accruing mutations in their germline VDJ genes (somatic hypermutation; SHM), as well as switching expression of constant region genes through DNA deletion events (class switch recombination; CSR). These endogenous processes have been used to make lineage inferences *in vivo*. We reasoned we could use these endogenous genetic alterations in the BCR as estimators of the initial cell state of any given clone detected *in vitro*. For example, we infer a B cell with an unmutated BCR detected in the time-course arose from a Naive B cell progenitor.

We assessed the validity of this approach by quantifying the concordance between transcriptionally defined memory and naive B cell states and categorically delineated SHM levels (germline, mutated, heavily mutated). In general, the concordance between mutational and transcriptionally-defined cell state categories was high (S2A, S2B). For example, in the Day 0 population more than 90% Naive B cells possessed an unmutated BCR gene, also known as a germline gene, and less than 3% of plasma cells were classified germline. In contrast, Non-Switched Memory B cells were more evenly split between germline and mutated (ratio 1.5 respectively), suggesting B cells can gain a transcriptional memory phenotype in the absence of detectable SHM. Critically, we observed no appreciable evidence of hypermutation *in vitro (*S2C), consistent with prior literature (Bergthorsdottir et al. 2001).

Given the germline and mutated categories specified Naive and Memory transcriptional states in the input population, we used the mutation level measured for *in vitro* differentiated B cells as a direct indication of whether their progenitor cell was a Naive or Memory B cell. We found the progeny of memory B cells increased two-fold in relative abundance over the course of the culture, showing memory B cells are intrinsically two-fold more persistent *in vitro (*2A). This is consistent with orthogonal measurements, which report memory B cells are on average one division ahead of Naive B cells when cultured *in vitro* (Tangye et al. 2003).

We continued to use the genetic status of the IgH locus as a lineage marker, which allowed us to understand the population dynamics of cell states *in vitro*. We performed differential gene expression analysis on the germline and mutated B cells, which identified distinct transcriptional programs amongst these differentiating subsets. Interestingly, we found mutated B cells were far more likely to express genes involved in T cell interaction (2B), suggesting Memory B cells are intrinsically licensed to enter an inflammatory state which activates T cells. In contrast, Naive B cells have a more limited ability to do the same. Instead, Naive B cells were biased toward expressing lectins and CCR7, suggesting Naive B cells are intrinsically primed to home into the lymphatic system and germinal centers (2B).

We quantified the relative magnitude of these cell fate biases (2C). At Day 0, we found what we expected in the peripheral blood, for example, that plasma cells were 4 times more often mutated than germline. However, by Day 4 germline cells populated the plasmablast/cell state almost as often as mutated cells, definitively linking Naive B cell progenitors to plasmablast phenotypes in this culture system. As the differentiation proceeded, the balance of mutated to germline B cells shifted back towards mutated, showing plasmablasts generated by mutated cells were more persistent *in vitro*.

Our measurement of phenotypes was not limited to the transcriptome because B cells generated additional phenotypic diversity *in vitro* through CSR. Generally, as Naive IGHM B cells experience cytokinetic/antigenic stimulation, they class-switch to any of the IGHA, IGHG or IGHE genes. This process diversifies the immune response by producing antibodies with the same specificity, but different effector functions. We quantified the *in vitro* dynamics of CSR through the lens of mutation status, which revealed strongly different fate biases between germline and mutated cells (2D). Most strikingly, B cells which switched to IGHE were almost exclusively derived from germline progenitors: the ratio of germline IGHE cells to mutated IGHE cells was (8-fold - inf, 95 % CI). We illustrate models of cell state and isotype biases in 2E.

### Clonal analysis reveals primed progenitors, clonal fate-bias, and map of class-switch events

We next used the full B cell receptor sequence to define clones in our dataset (Methods). Among the 11,171 differentiating B cells, 1,911 were related to at least one other detected cell (3A, S3A). We measured the cell states of many clonal families, and determined that clones had strong fate biases; clones were three-fold more likely to be found in a single cell fate than a null expectation (3B). Amongst the clonal families, we detected 73 families whose members were detected at Day 0 and at a later time point. We called these persistent clones, and observed Non-Switched Memory B cells appeared to be the most common cell state at Day 0 in persistent clones. This suggests Non-Switched Memory B cells were better at proliferating in these conditions. To test this hypothesis, we modeled a scenario in which persistence was even amongst all Day 0 clones: division rates were the same, and death rates were zero. We detected ∼2X more progeny of Non-Switched Memory B cells than would be expected in the case of the aforementioned even–expansion model, indicating strong cell intrinsic biases toward *in vitro* persistence amongst progenitor cells (3C).

We also used the clonal information to understand the *in vitro* dynamics of class-switch recombination. On the population level, we observed as much as an order of magnitude increase in the amount of class-switched cells above the input (S3C). We used the observed intra-clonal isotype counts to derive a map of class-switch outcomes *in vitro* (3D). For comparison, we calculated the naive probability of detecting a switch given the proportions of isotype usage in the general population. For same-same isotype relationships (i.e. IGHG1 IGHG1), the *in vitro* map of class-switching showed more than ten-fold enrichment compared to the naive probability model. This enrichment can be explained by clonal inheritance of isotype status. We also noted a strong divergence from this model for the IGHM to IGHA1 switch, where the naive probability model expects intraclonal switches which were not observed. This discrepancy strongly indicates the IGHA+ cells detected in later time points were derived from the persistence of IGHA cells already present in the input. Thus, CSR from IGHM cells did not meaningfully contribute to the abundance of IGHA+ cells in the population. In contrast, we noted that many intraclonal class-switching events appeared to be directly from IGHM to IGHE. Explanations involving unobserved cells with intermediate isotypes notwithstanding, these data illustrate the relative ease with which B cells can switch directly to IGHE. Our data for IGHE cells contrasts with *in vivo* data which show IgE B cells to be: (1) very rare, (2) apparently derived from sequential switching (e.g. from IgG1 to IgE) (Horns et al., 2016; Looney et al., 2016), and (3) often heavily hypermutated (Croote et al., 2018). Taken together, these data suggest that while direct switching to IGHE from Naive progenitors is trivial *in vitro*, niche factors or intrinsic death programs efficiently limit their generation or lifetime *in vivo*.

### Persistence of transcriptional memory varies across genetic loci

The intrinsic biases in fate outcomes that we detected must be underpinned by the persistence of transcriptional memory at individual genetic loci. In principle, some genes may exhibit faithful transmission of transcriptional state across clonal expansion, while other genes may not. Thus, we sought to determine how the persistence of transcriptional memory varies across the genome. We employed a permutation test on the clone labels to find genes which were less variable within clones than between. Using this test, we identified a set of 6,937 genes with p-values less than < 0.01. Supporting the sensitivity of our test, it identified genes known to be clonally inherited: the light chain variable genes and the light chain constant regions. These genes were not used *per se* to identify clones, providing evidence for the validity of the test. We call the genes we identified clonally coherent genes (CCGs). One way to intuitively understand these clonal effects is to observe the cascade plot of the putative CCG, for example IGKC (4A) (Cheon et al., 2021). A clear clonal structure of high expressing clones and low expressing clones is scrambled upon permuting clonal labels. Whether we detected a CCG was highly related to its expression level. Most highly expressed genes were clonal, and unsurprisingly we often could not reject the null hypothesis for genes which were difficult to detect (4C).

We calculated an effect size of the variability in gene expression explained by the clonal labels called the clonal index. Unsurprisingly, the Ig variable genes and light chain genes had some of the strongest effects (4B). In contrast to the light chain, where cells generally only transcribed one constant region gene, we measured robust transcription from multiple different heavy-chain constant region genes in single cells (S4C). This transcription is consistent with the so-called sterile transcription which is necessary for CSR (Lee et al., 2001). Strikingly, we found Ig heavy-chain constant region genes were CCGs, which was surprising given the diversity of transcriptional states measured for the locus. The clonal coherence of IgH transcription suggests faithful propagation of chromatin states at the IgH locus across cell division, and may be an explanation for observations of clonal coherence in isotype usage *in vivo (Horns et al., 2016)*. We noted that the CCGs with the strongest effects were often genes known B cell fates, such as MS4A1, AIDCA, and JCHAIN. To quantify these observations, we calculated the overlap coefficient between the set of top CCGs and top Differentially Expressed Genes (DEGs) between B cell states. We observed strong agreement between the set of top DEGs and top CCGs (0.39 overlap), which was fourteen-fold higher than the agreement between DEGs and null-sets of genes sampled from the B cell transcriptome. Using a set of important B cell genes curated from the literature (Morgan and Tergaonkar 2022), we also found an 8-fold enrichment for CCGs. These enrichments indicate the CCGs are involved in cell fate determination and are relevant features for functional characterization.

Finally, we asked whether we could identify CCGs from *in vivo* samples. To this end, we performed the permutation test on only the clones in our long-lived plasma cells from the bone marrow. We found an order of magnitude fewer genes were detected as clonal and, of these, 561 genes were non-Ig genes (S4B). Nonetheless, the genes we identified were important for plasma cell function such as MZB1, JCHAIN, and KRTCAP2. JCHAIN was detected as a CCG both *in vitro* and *in vivo*. Thus, we speculate that differentiating B cells make a stable fate choice to secrete polymeric or monomeric Ig. Given the plausible age of a CD38+CD138+ plasma cell (Hammarlund et al., 2017), we expect we are detecting inherited transcriptional programs on the scale of years later.

## Discussion

An effective immune response requires the profound and rapid activation of phenotypic diversity while maintaining tight control over proliferation and inflammation. We combined single-cell genomics and lineage tracing to investigate the activation process of human primary B cells. While we observed a diversity of phenotypes in the population of activated B cells, we were able to explain much of this diversity via inference of progenitor states via lineage tracing. For example, memory B cells were more persistent in culture and had a stronger license to present pro-inflammatory factors. Meanwhile, Naive B cells almost entirely accounted for the IGHE cells, likely indicating their relative sensitivity to IL4 stimulation. Our clonal analysis detected strong cell fate biases, which are likely underpinned by the CCGs we identified. We detected CCGs in terminally differentiated plasma cells from an *in vivo* sample, revealing clonal transcriptional programs may be stable for years *in vivo*. The set of CCGs was heavily enriched for cell-type defining genes such as MZB1, JCHAIN, and IGHE, while still containing hundreds of genes which are not yet described in B cell biology.

Our results illustrate how lineage tracing in a cell reprogramming protocol can provide fruitful insight into the intrinsic biases of progenitor cells, which will ultimately allow better control over cell reprogramming outcomes. *In vitro* differentiation is not just common in research labs, but is now clinically useful in the form of CAR-T therapy and regenerative medicine efforts. Single cell genomics already offers powerful insight into cell therapeutics (Wang et al. 2021; Bai et al. 2022), but adding *in vivo* and *in vitro* lineage tracing technologies will yield critical additional information about how to improve targeting and potency. We also expect the clonal effects and intrinsic biases identified here play a powerful role in biasing cells towards particular effector fates *in vivo*.

Here we exploited the unique biology of B cells to gain insight into their differentiation processes. For example, we conceptualized the BCR an *in vivo* molecular recorder of germinal center experience. This allowed us to dissect the striking differences in activation programs between memory and naive B cells. These differences are particularly interesting given these cell types are transcriptomically difficult to distinguish when at rest. In general, as lineage tracing and cell-recording technologies continue to develop, researchers will genetically record bespoke cell experiences, and use single-cell genomics to analyze their effects on intrinsic cell biases (Chen et al., 2022). Importantly, the intrinsic differences between B cell clones became most apparent when we subjected them to a differentiation protocol *in vitro*. As single cell genomics atlases broaden our understanding of human tissue biology (Eraslan et al., 2022; Tabula Sapiens Consortium* et al., 2022), it may be critical to perform *in vitro* perturbations on these tissues to understand the full diversity of human biology.

## Supplementary Figures

**Figure S1.**
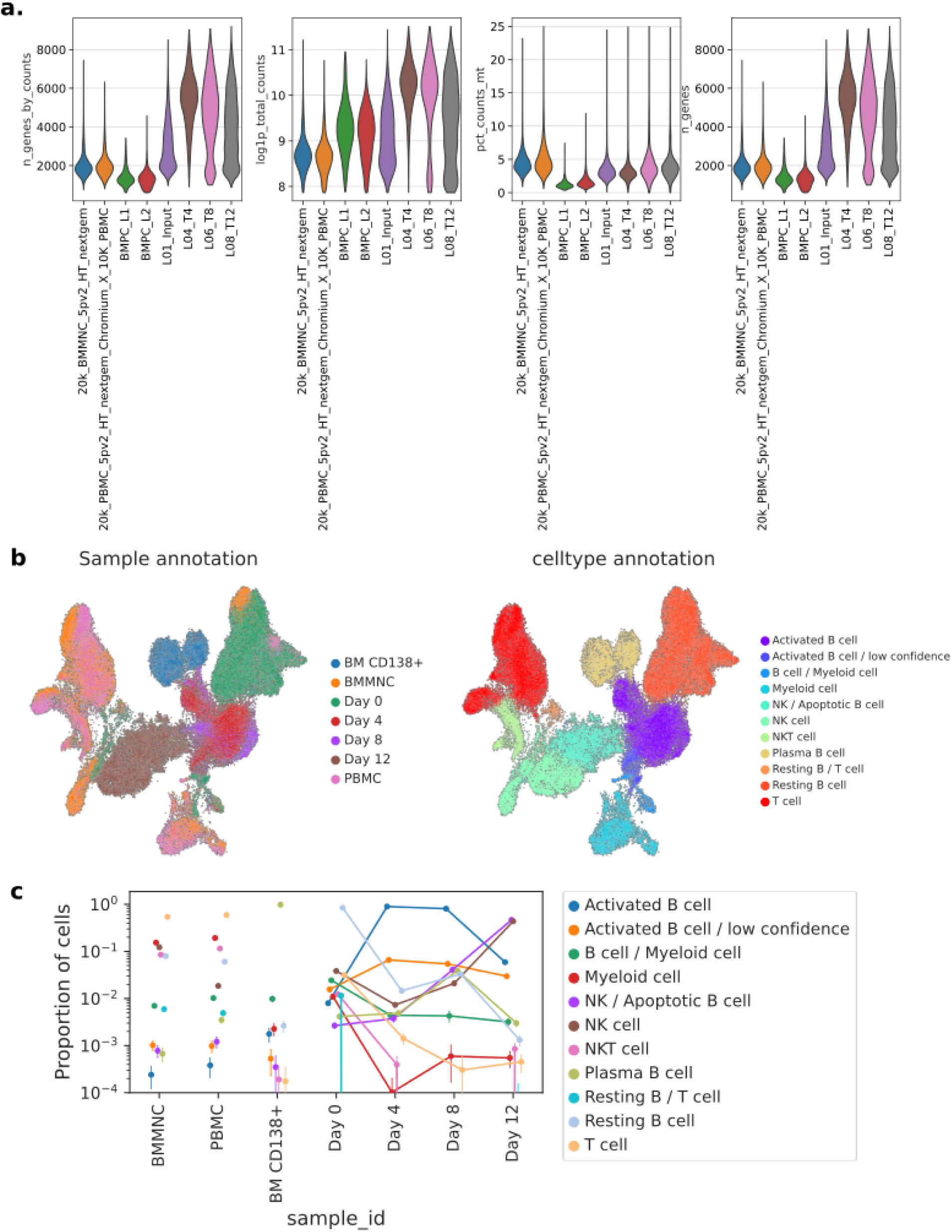
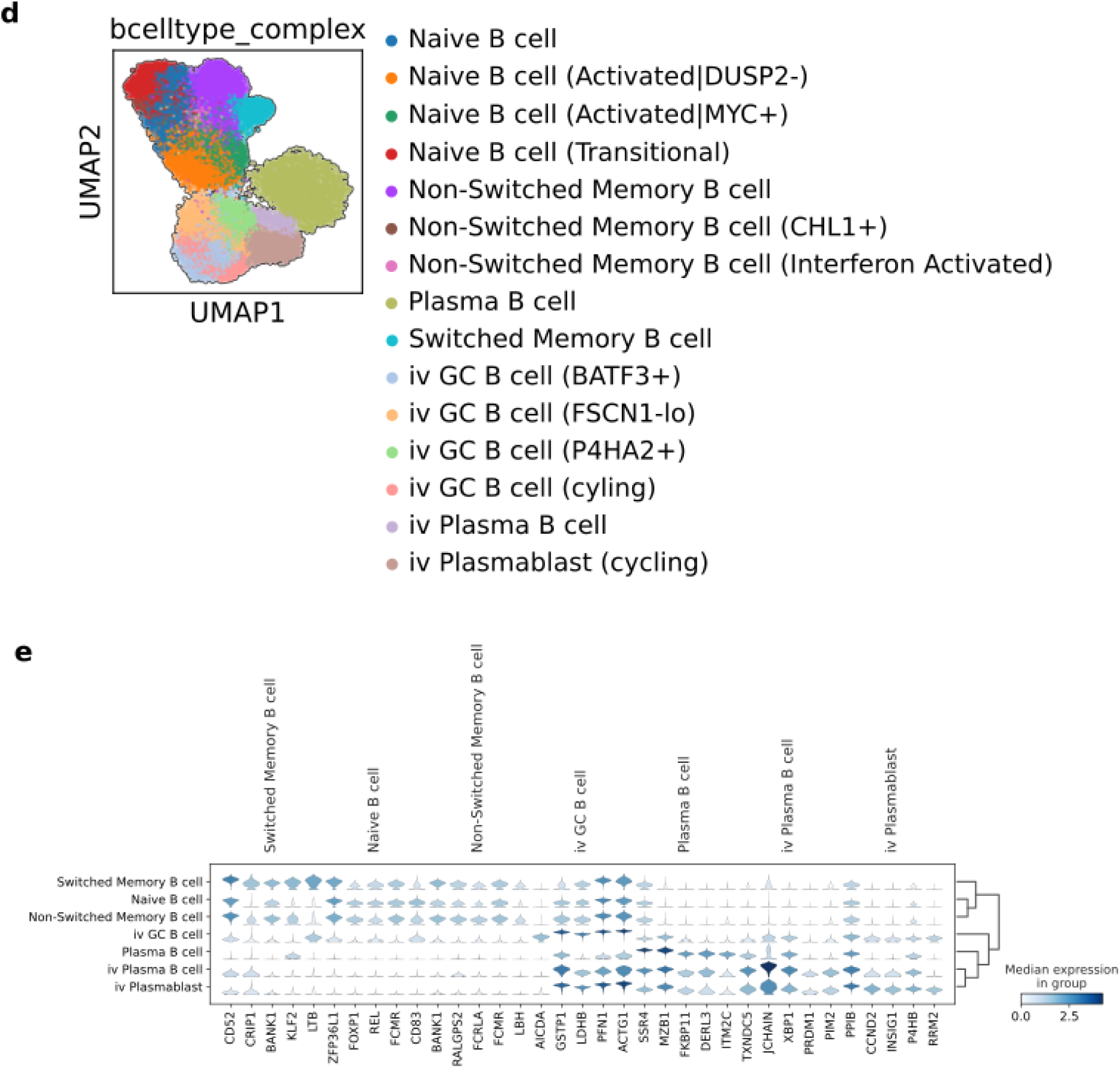
(A) Violin plots of quality metrics for each 10X genomics lane sequenced and/or analyzed (B) UMAPs of all cells in the dataset, colored as shown in legends. Contaminant cells in the B cell purifications cluster with the similar cells in unpurified fractions (C) Pointplot quantifying the proportion of celltypes in each sample_id. (D) Sub Clustering of B cell type annotations with greater leiden resolution. These higher resolution “fates” were used for the histogram in 3B (E) Differentially expressed genes which define each B cell type.

**Figure S2.**
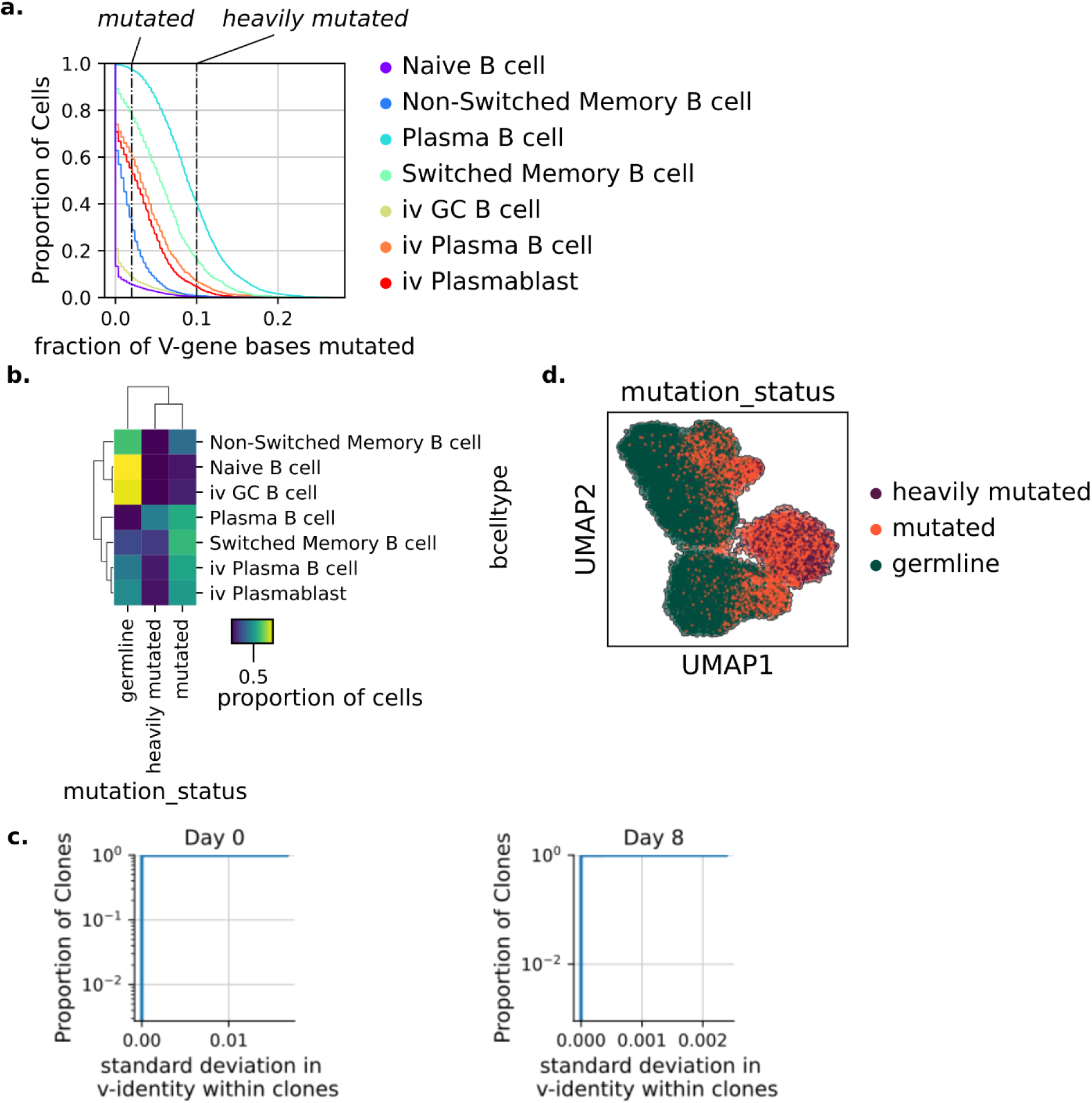
(A) Empirical cumulative distributions of the fraction of V-gene based mutated away from the germline V-gene, for each transcriptomically defined B cell type (B) A confusion matrix showing the concordance between the mutation status based on S2A and each B cell type label based on leiden clustering. (C) Empirical cumulative distributions of the standard deviation in v-identity within clones shows mutations are not collected within clones over the time course (D) UMAP plot colored by the mutation status assigned based on S2A.

**Figure S3.**
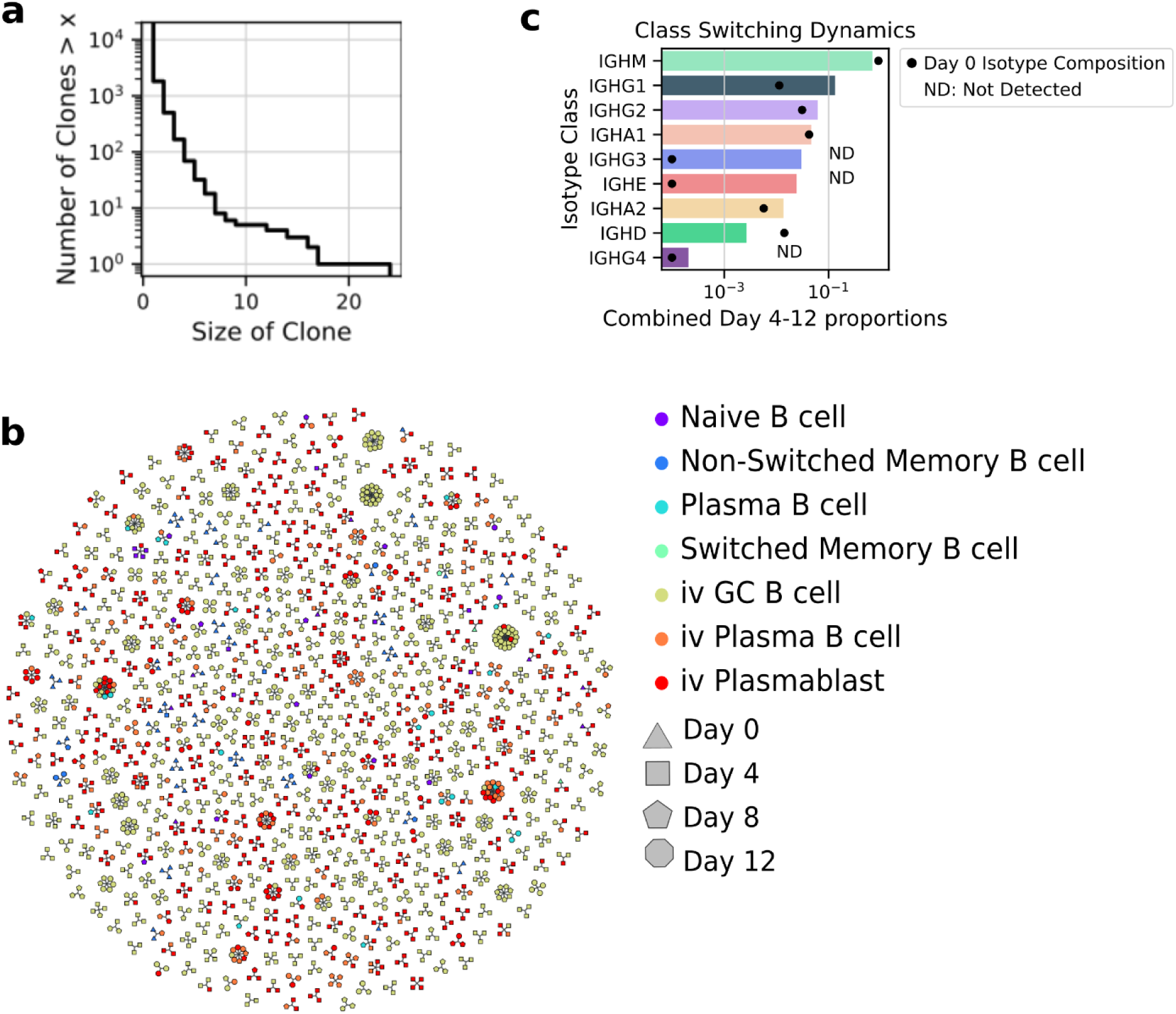
(A) Clone size distribution of the B cell population from the *in vitro* time-course (B) A graph based representation of the persistent clones: clones which were detected in multiple time points. (C) Class-switching (isotype-switching) dynamics during the culture. ND = Not Detected in the Day 0 population, a pseudo-count is added for visualization purposes.

**Figure S4.**
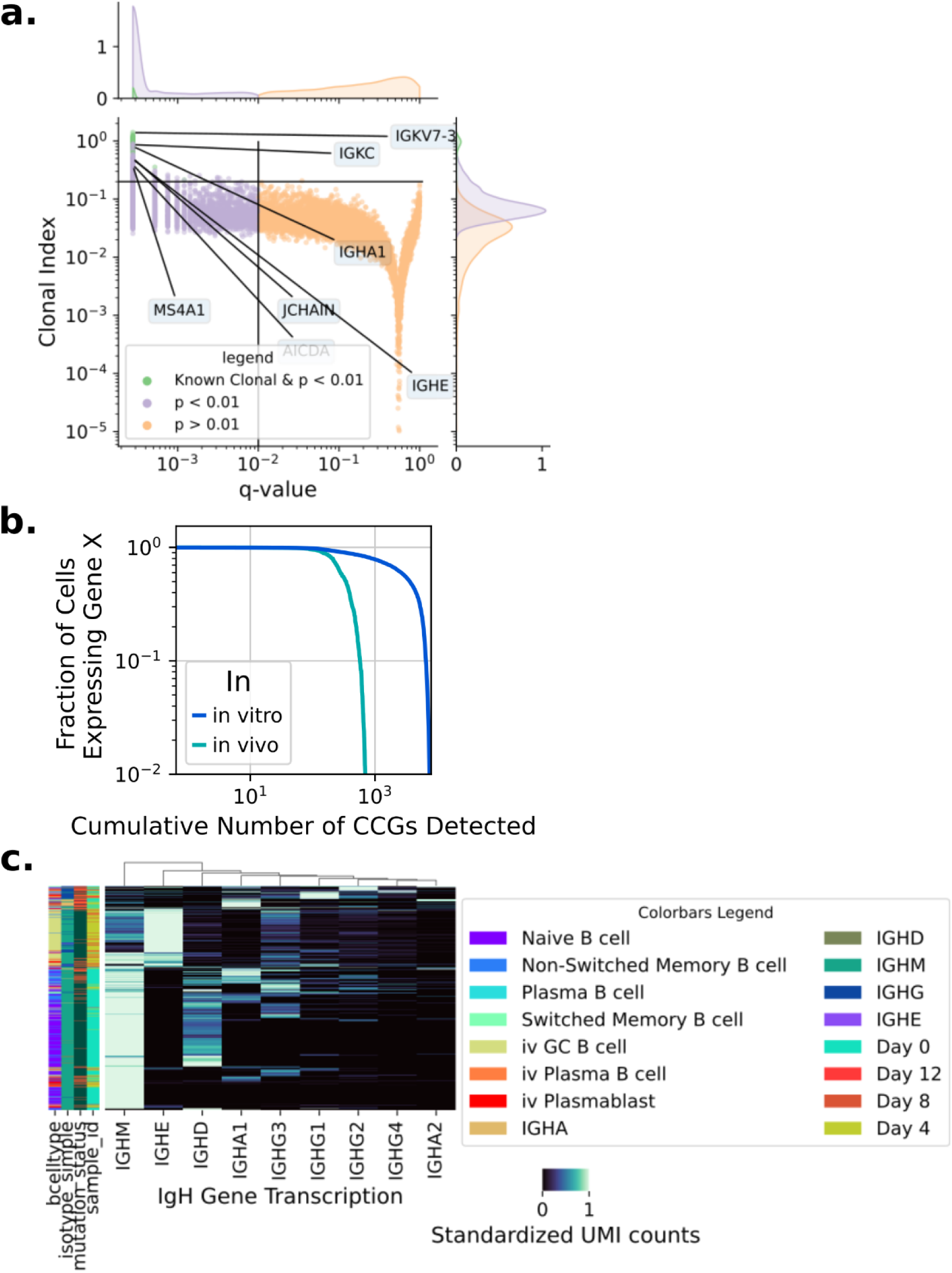
(A) A volcano plot showing results of the transcriptome-wide permutation test only for the *in vivo* (BM CD138+) sample. The q-values are Benjamini-Hochberg corrected p-values and the clonal index is a normalized metric of expression variance described in the methods. Genes of interest are labeled and groups of interest are colored. (B) Empirical cumulative distribution plot of the number of clonal genes detected with q-value < 0.01 by the Fraction of cells expressing a given gene (also known as gene drop-out). (C) Clustermap of cells by transcripts detected at the IgH locus, standardized within each row (cell)

## Materials and Methods

### Sample collection

PBMCs were obtained from 2 healthy adult male LRS chambers from the Stanford Blood Center. LRS chamber product was diluted 1:4 in PBS + 2% FBS and PBMCs were isolated using a Ficoll gradient and Red Blood Cell Lysis. Cells were frozen in Cryostor CS10 according to the manufacturer’s instructions. Human bone marrow aspirates from a healthy male aged 50-55 were obtained from AllCells. Mononuclear cells were isolated using Ficoll gradient and Red Blood Cell Lysis. Plasma cells from the bone marrow were isolated using StemCell EasySepEasySep™ Human CD138 Positive Selection Kit II

### Experimental Model and Subject Details

Study subjects gave informed consent and protocols were approved by the Stanford Institutional Review Board and/or the AllCells Institutional Review Board.

### B cell purification and culture

PBMCs were thawed and B cells were purified by negative selection using the StemCell B cell enrichment kit. The resulting cells were measured by flow cytometry to be over 84% pure, which was confirmed by single-cell RNA sequencing. B cells were cultured for 8 hours in B cell media (StemCell) at 37 C 5% CO2 at a density of 2 × 10^4^ cells per well. After 8 hours B cells were stimulated using the B cell stimulation cocktail according to manufacturer’s instructions (Stem Cell). B cell culture wells were thoroughly mixed every 24 hours, to mitigate any spatial effects of the culture on particular B cells. On Day 4, the cells were split into 3 wells and restimulated. Every 24 hours these three wells were pooled, mixed and redistributed into three new wells. On Day 8, the B cells were restimulated and separated into 6 wells, at which point the pooling and mixing was carried out using 6 wells.

### Single-cell isolation and sequencing

B cells were washed 2 times in PBS + 1% BSA and stained with cite-seq antibodies according to (Stoeckius et al., 2018). Then, cells were counted and loaded on the Chromium (10X Genomics) at a target loading of 20,000 cells per lane. Reverse transcription and complementary DNA (cDNA) amplification were performed using the Single Cell V(D)J kit V2 (10X Genomics). VDJ and gene expression libraries were prepared from each of the 4 time points. All library preparation was done according to manufacturer’s instructions, except for the use of custom constant region primers (Horns et al., 2016) for VDJ enrichment. On Days 8, and 12, cells from culture wells were sorted by propidium iodide exclusion into cold PBS + 2% BSA, before proceeding to antibody staining. Libraries were sequenced using the Illumina NovaSeq platform with paired-end reads of 26 bp and 98 bp.

### All analysis code

https://github.com/michael-swift/seqclone3

### Preprocessing of single-cell sequence data

We used snakemake (Mölder et al., 2021) to manage the computational workflow. We used CellRanger 6.1 to map, count, and assemble reads from the sequencing libraries. We used the Immcantation docker (Gupta et al., 2015) pipeline and scirpy to reprocess the contigs assembled by CellRanger, distinguish bonafide single cells from multiplets, and assign clonal barcodes, which agreed with our in-house pipeline (Croote et al., 2018). Single B cells were identified by the presence of a single productive heavy chain and a single productive light chain, yielding a total of 29,703 single B cells for analysis. All other cells were excluded from further analysis.

### Single cell sequencing data analysis

Gene expression analysis of single cells was performed using scanpy (Wolf et al., 2018), and exploratory analysis of the immune receptors with scirpy (Sturm et al., 2020). Briefly, single cell transcriptomes were log-transformed and normalized to counts per 10^4^ UMIs. Clusters were identified using the leiden algorithm. These clusters were manually annotated (i.e. labels were assigned) based on expression of marker genes for each cell state. Differentially expressed genes were identified using the wilcoxon-rank sum test adjusted for multiple testing using the Benjamini-Hochberg procedure as implemented in scanpy.

### Analysis of mutated vs. germline outcomes

To explore the cell fate biases of memory vs. naive progenitors, we binned VDJ sequences into heavily mutated, mutated, and germline categories as shown in S2A. We used multiple different levels of reasonable cutoffs and found they did not change our general conclusions. We calculated the ratio of germline over mutated by dividing the number of germline cells by the number of mutated or heavily mutated cells in a given type of category, such as which IGH constant region is associated with the cell’s VDJ or which b cell type is associated with a VDJ.

### Analysis of clonal gene expression

In general we employed the resampling and permutation methods to estimate confidence intervals and p-values for all clonal effects. We implemented the approach lucidly described here (Horton et al., 2018), and all code implementing these tests will be publicly available on GitHub. The test statistic for genes was the within-clone variance in UMI counts for each gene averaged across all clones. For calculating the clonal index in 4B, we divided the difference in the gene-statistics between true labels and permuted labels by mean transformed UMI count for that gene.

## Acknowledgements

We thank Ivana Cvijovic, Elisabeth Jerison, Derek Croote, Bali Pulendran, Dan Jarosz, and Daria-Mochly Rosen for useful discussions and helpful comments on the manuscript. This work was supported by the National Science Foundation Graduate Research Fellowship Program (to M.A.S.)

