## Supplementary Files for "Fate Bias and Transcriptional Memory of human B cells"

### Supplementary Material:

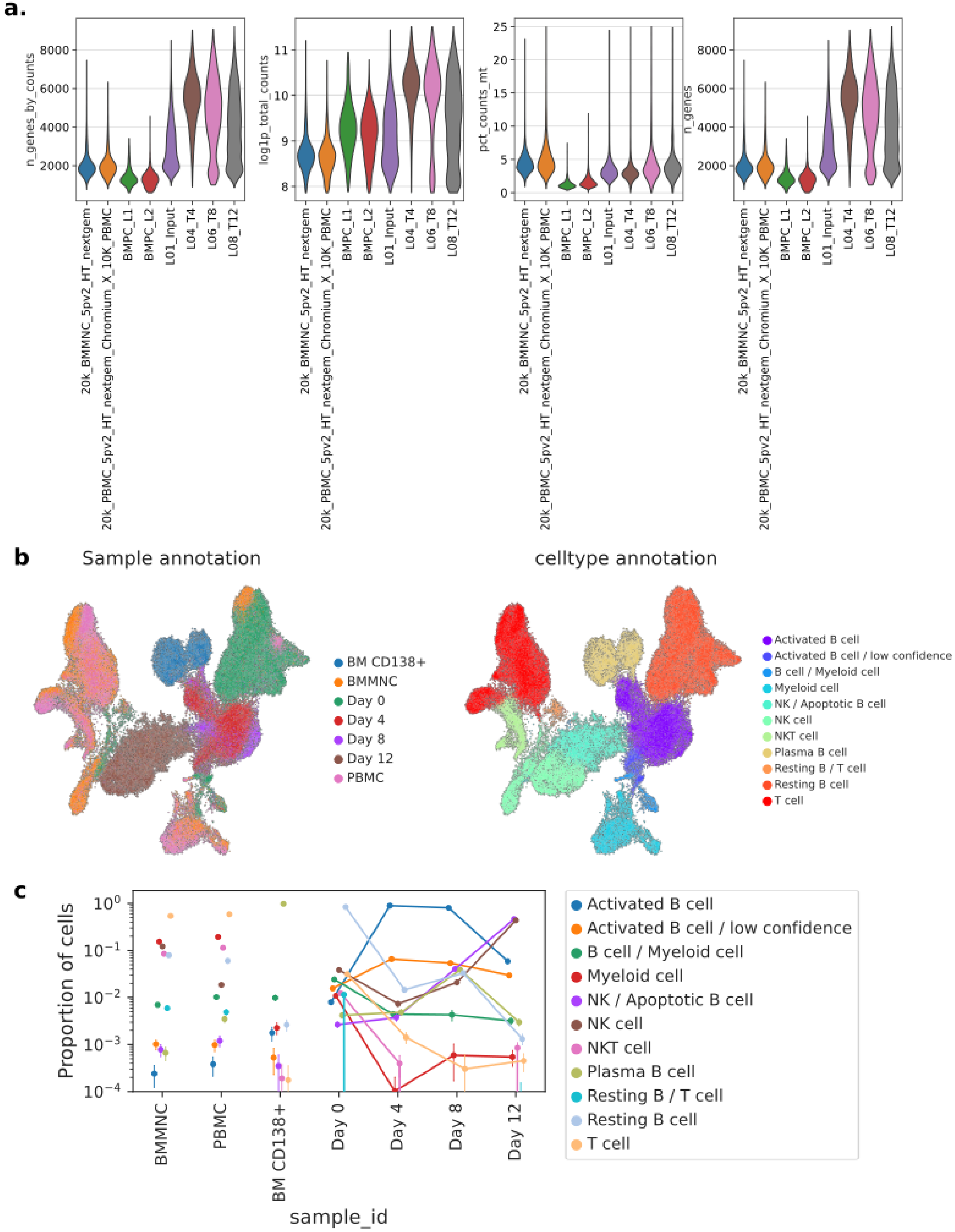
a.
n_genes_by_counts
log1p_total_counts
pct_counts_mt
n_genes
20k_BMMNC_5pv2_HT_nextgem
20k_PBMC_5pv2_HT_nextgem_Chromium_X_10K_PBMC
BMPC_L1
BMPC_L2
L01_Input
L04_T4
L06_T8
L08_T12
b
Sample annotation
celltype annotation
BM CD138+
BMMNC
Day 0
Day 4
Day 8
Day 12
PBMC
Activated B cell
Activated B cell / low confidence
B cell / Myeloid cell
Myeloid cell
NK / Apoptotic B cell
NK cell
NKT cell
Plasma B cell
Resting B / T cell
Resting B cell
T cell
c
Proportion of cells
sample_id
BMMNC
PBMC
BM CD138+
Day 0
Day 4
Day 8
Day 12

d

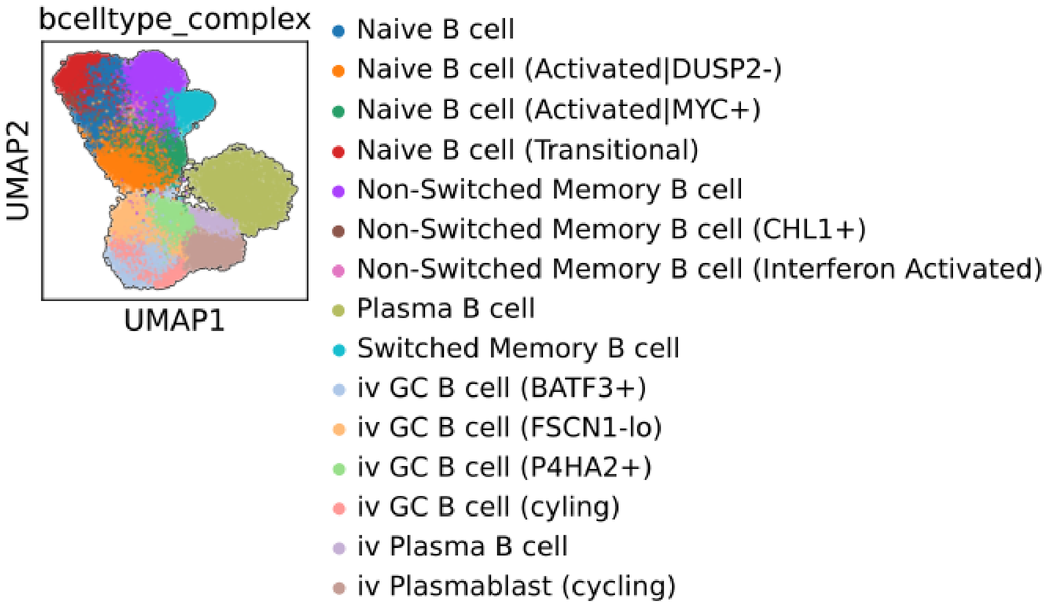


e

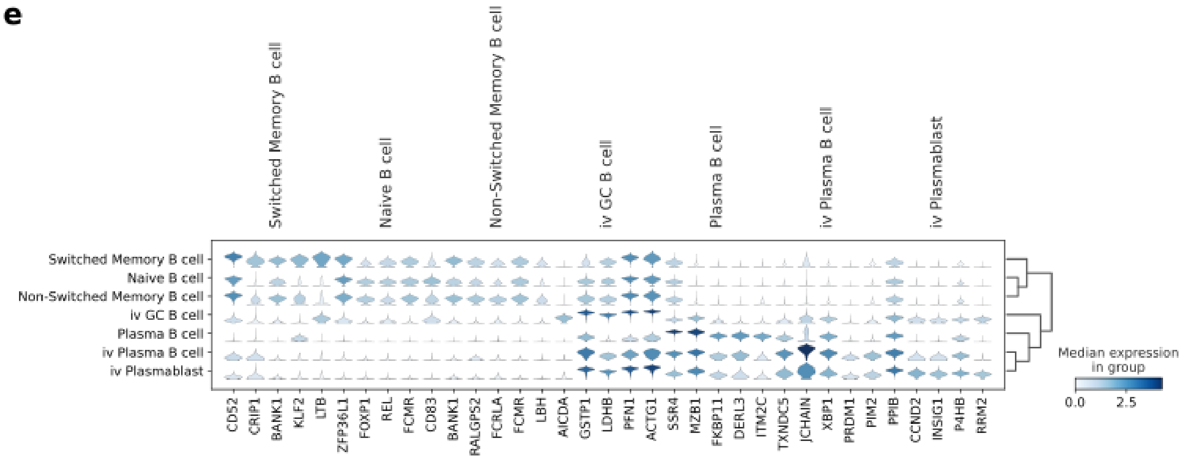


**Figure S1.** (A) Violin plots of quality metrics for each 10X genomics lane sequenced and/or analyzed (B) UMAPs of all cells in the dataset, colored as shown in legends. Contaminant cells in the B cell purifications cluster with the similar cells in unpurified fractions (C) Pointplot quantifying the proportion of celltypes in each sample_id. (D) Subclustering of B cell type annotations with greater leiden resolution. (E) Differentially expressed genes which define each B cell type.

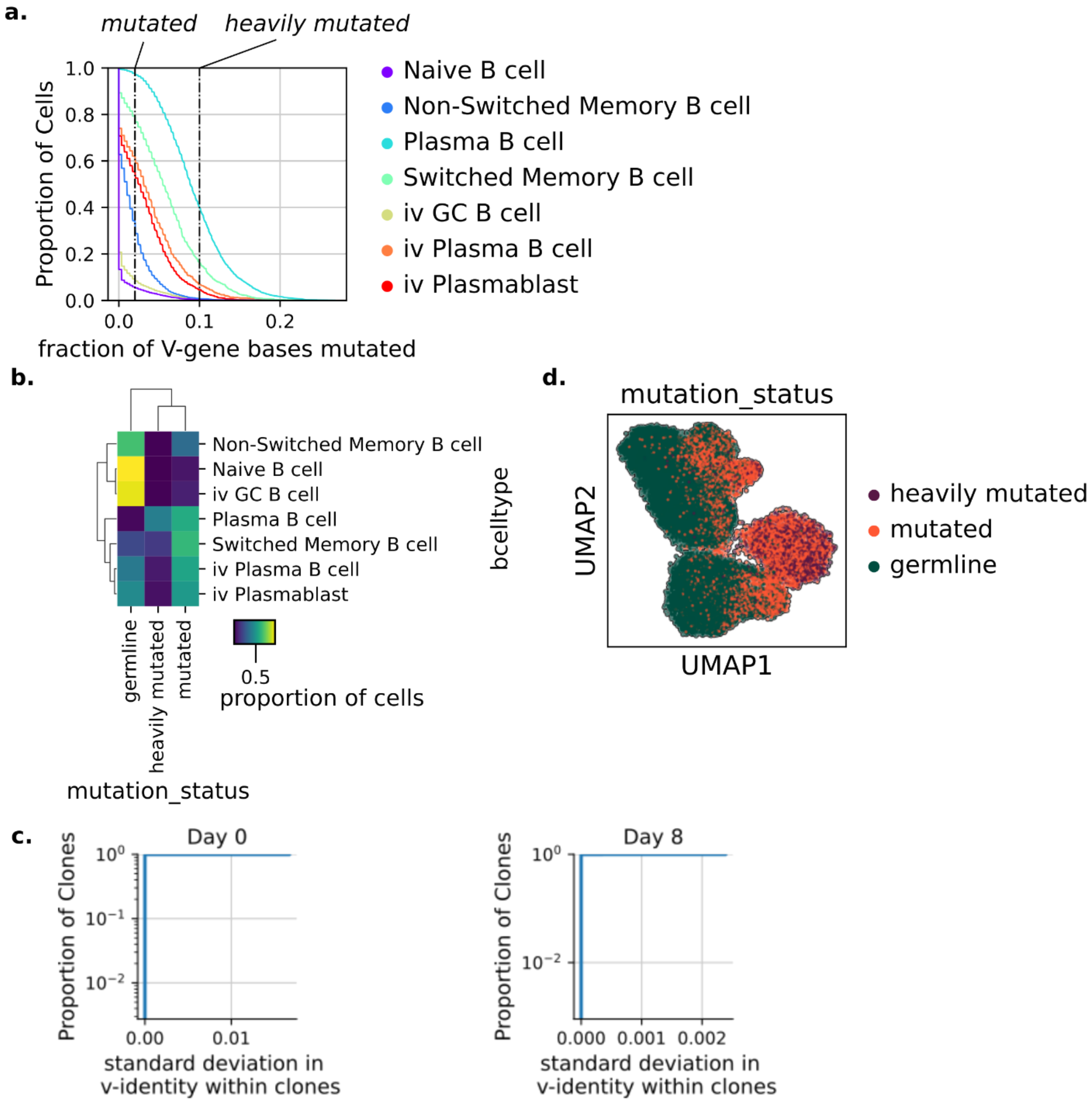


**Figure S2.** (A) Empirical cumulative distributions of the fraction of V-gene based mutated away from the germline V-gene, for each transcriptomically defined B cell type (B) A confusion matrix showing the concordance between the mutation status based on S2A and each B cell type label based on leiden clustering. (C) Empirical cumulative distributions of the standard deviation in v-identity within clones shows mutations are not collected within clones over the time course (D) UMAP plot colored by the mutation status assigned based on S2A.

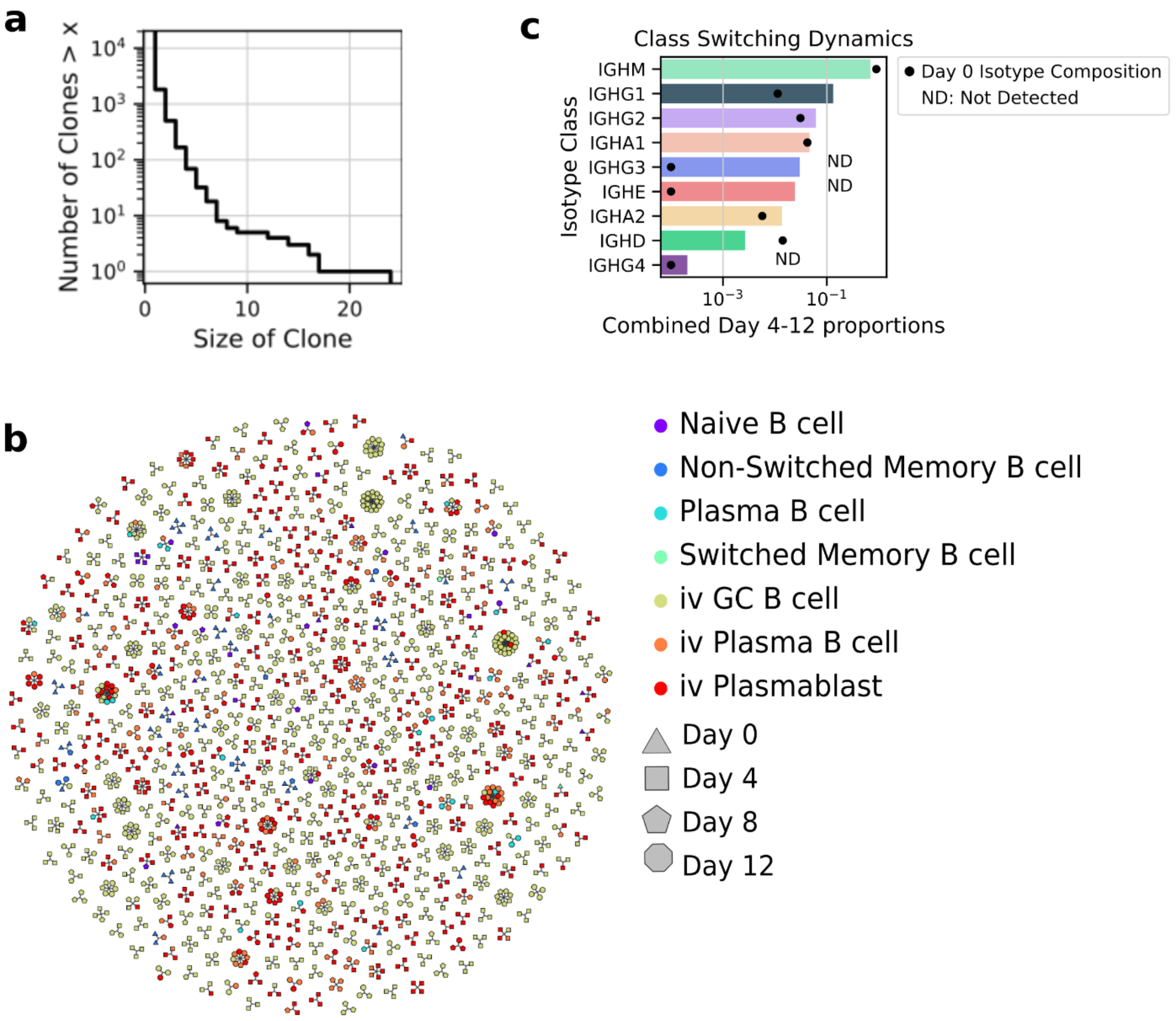


**Figure S3.** (A) Clone size distribution of the B cell population from the *in vitro* time-course (B) A graph based representation of the persistent clones: clones which were detected in multiple time points. © Class-switching (isotype-switching) dynamics during the culture. ND = Not Detected in the Day 0 population, a pseudo-count is added for visualization purposes.

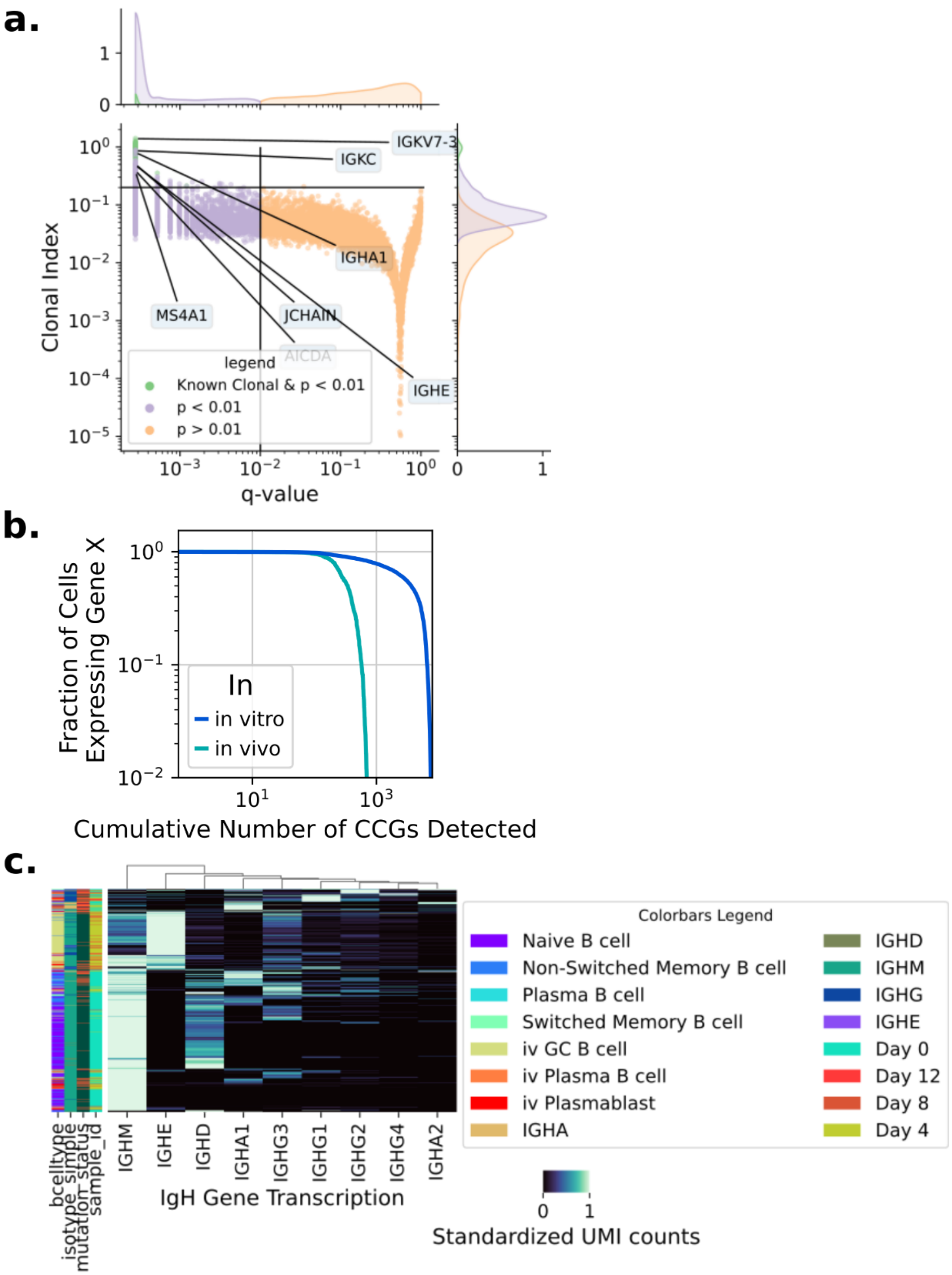


**Figure S4.** (A) A volcano plot showing results of the transcriptome-wide permutation test only for the *in vivo* (BM CD138+) sample. The q-values are Benjamini-Hochberg corrected p-values and the clonal index is a normalized metric of expression variance described in the methods. Genes of interest are labeled and groups of interest are colored. (B) Empirical cumulative distribution plot of the number of clonal genes detected with q-value < 0.01 by the Fraction of cells expressing a given gene (also known as gene drop-out). (C) Clustermap of cells by transcripts detected at the IgH locus, standardized within each row (cell)
